# Phosphorus limitation enhances root exudation and mineral bioweathering across diverse soil process domains

**DOI:** 10.64898/2026.02.13.705823

**Authors:** Sasha Pollet, Jean-Thomas Cornelis, Thorsten Knipfer, Cindy Prescott, Kylee Tate, Young-Mo Kim, Guillaume Lobet

## Abstract

**Aims:** Harnessing rhizosphere processes offers a valuable opportunity to optimize nutrient use efficiency in agroecosystems. In nutrient-limited soils, plants discharge part of photosynthate surplus via root exudation, including carboxylates, which may enhance mineral dissolution and nutrient mobilization. We aimed to assess how plant responses to nutrient limitation translated into changes in exudate profiles, and how these exudates, in turn, drive bioweathering across soils of contrasting mineralogy and weathering degree.

**Methods:** We conducted a hydroponic experiment with *Lupinus albus* grown in a phosphorus (P) gradient over seven weeks. We measured plant biomass and root traits, performed a metabolomics analysis and quantified seven carboxylates in root exudates using gas chromatography-mass spectrometry. To assess bioweathering across contrasted soil domains, we conducted batch dissolution tests with exudates using three soil horizons—each with distinct physicochemical properties: enriched in organic matter, iron oxides, or primary silicates.

**Results:** At the intermediate level of P supply, shoot biomass was comparable to that under high P, but plants produced more root biomass and a higher total carboxylate exudation rate. Despite low carboxylate concentrations (<100 ppb), exudates promoted the dissolution of Ca, Mg, Si, Fe, P and K in all soils. Yet, the degree of element released varied among soil types.

**Conclusion:** These findings highlight the importance of root exudates in enhancing mineral dissolution, with effects dependent on soil physicochemical properties. The results suggest that managing agroecosystems under moderate nutrient limitation could be a sustainable strategy to increase root-to-shoot ratios, enhance bioweathering and nutrient release in rhizosphere.

## Introduction

The Green Revolution of the twentieth century, which brought tremendous gains in crop yield, was largely driven by the use of mineral fertilizers and plant breeding (Pingali, 2012). Agriculture intensification based on external inputs has contributed to pollution and global warming, raising significant concerns about the ecological footprint of agriculture (Smith et al., 2016). To attain the twin goals of feeding the world’s population while keeping Earth’s system in balance (Steffen et al., 2015), we need a paradigm shift which will harness soil-root-plant interactions benefits to build resilience and sustainability in agroecosystems (Bishopp & Lynch, 2015; J. P. Lynch, 2007; J. P. Lynch et al., 2022). Harnessing rhizosphere processes presents an opportunity to optimize nutrient use efficiency in agroecosystems (Oburger et al., 2022; Preece & Peñuelas, 2020; Schnepf et al., 2022; C. Wang & Kuzyakov, 2023).

Plants are major drivers of rock weathering (Dontsova et al., 2020; Porder, 2019). They can physically or chemically weather soil minerals, which release elements in rhizosphere that can be taken up by plants and microorganisms. In natural systems, bioweathering rates (i.e. the biological processes of weathering that break down minerals) are maximized in nutrient-impoverished soils, where over millions of years, plants and their symbionts have developed strategies to cope with nutrient limitations (Lambers et al., 2008; Teste & Laliberté, 2019; Zemunik et al., 2015). By capitalizing on natural plant strategies, managing cropping systems under optimized nutrient deficiency is a promising way to promote plant nutrient acquisition and microbial C supply, without unduly compromising plant aboveground biomass (Prescott et al., 2021; C. Wang et al., 2025). Under sub-optimal nutrient conditions, leaf growth is curtailed more than carbon (C) fixation (Assuero et al., 2004; Kavanová et al., 2006), creating a surplus of photosynthates that are exported from the leaf, into multiple sink organs. Different nutrient limitations trigger distinct cascade mechanisms of plant and root responses, leading to changes in the metabolite profile of root exudates (Carvalhais et al., 2011), and stimulating rhizospheric activity (Francis et al., 2023; Freschet et al., 2021a). During soil genesis, the type of nutrient limitation changes (Lambers et al., 2008). In recently-formed soils, primary production tends to be N-limited, whereas in ancient soils, P is the limiting resource for plants, as P reserves in the minerals have been depleted (Vitousek et al., 2010). As root exudate composition is influenced by the nutrients that are limiting plant growth (Carvalhais et al., 2011; Jones, 1998), changes in nutrient limitation, weathering degree and mineralogy during soil genesis are expected to drive shift in exudates profile. Yet, how root exudation responds to nutrient limitation over soil genesis remains poorly characterized.

Root exudates are comprised of primary and secondary metabolites of both low molecular weight (MW;<1000 Da; e.g. sugars, organic acids, phenolics, vitamins) and high MW (>1000 Da; e.g. enzymes, mucilage) (Oburger & Jones, 2018). They constitute up to 20% of plant fixed C (Lynch & Whipps, 1990) and play a critical role in biogeochemical processes and ecosystem responses to environmental change (Li et al., 2021; Ma et al., 2022; Panchal et al., 2022; Shabtai et al., 2024). Root exudates play a pivotal role in plant functioning through nutrient acquisition by direct nutrient mobilization or via interactions with symbiotic fungi (Sardans et al., 2023, Dontsova et al., 2020; Porder, 2019; Wild et al., 2022). Key biotic factors influencing root exudation profiles include plant genotypes, developmental stage and interaction with rhizosphere microbes (McLaughlin et al., 2023; Naveed et al., 2017). In addition to soil chemistry, abiotic factors such as drought, high salinity, atmosphere CO_2_ concentration, and temperature also influence root exudation (Vives-Peris et al., 2020). Some exuded metabolites contribute uniquely to rhizosphere dynamics—for example, carboxylate efflux plays a crucial role in plants’ acquisition of phosphorus (P), manganese (Mn), and silicon (Si), as well as crops’ tolerance to aluminum (Al) (Dakora & Phillips, 2002; de Tombeur et al., 2021; Gerke, 2018; Ofoe et al., 2023). Phenolic and phytosiderophores contribute to iron (Fe) mobilization (Chai & Schachtman, 2022) and secondary metabolites such as flavonoids and strigolactones facilitate plant associations with mycorrhizal fungi and N-fixing symbionts, particularly under drought and P-deficient conditions (Hassan & Mathesius, 2012; Yoneyama et al., 2012). The effect of root exudates on element mobilization depends on the type of root exudates and the type of soil minerals (Bölscher et al., 2025; Liang et al., 2023; Veneklaas et al., 2003). Yet, the role of soil genesis—through its effect on soil mineralogy, element speciation, pH and organic matter dynamics—in shaping these interactions remains largely unexplored (Cornelis & de Tombeur, 2022).

Among root exudates, organic acids (OAs) and their conjugated based, carboxylates, have been studied for their effects on bioweathering processes (Cornelis & de Tombeur, 2022; Dakora & Phillips, 2002; Jones, 1998; Jones & Darrah, 1994; Oburger et al., 2009). Carboxylates impact bioweathering through 1) ligand-promoted dissolution (chelating molecules can promote dissolution of primary minerals by forming inner-sphere complexes at mineral surface; (Wild et al., 2022), 2) proton-promoted dissolution (at soil pH over 3-4, carboxylates are deprotonated, releasing protons in soil solution) and 3) affecting reaction affinity through organic-metal chelation of metal ions in the soil solution (Lawrence et al., 2014). Carboxylates have also been studied in P desorption processes as P is largely unavailable to plants due to adsorption on oxides and precipitation with metal cations (Ca in alkaline soil and Al, Fe in acidic soils) (Lambers, 2022). As nutrient-limited conditions alter plant stoichiometry and trigger cascading effects within plant systems (Göran I., 2008; Prescott et al., 2020), it is reasonable to expect shifts in root exudate profiles at different stages of soil genesis. How these altered root exudate profiles influence bioweathering processes across soil genesis remains, however, largely unknown.

*Lupinus albus* (white lupin) is a model species for studying adaptations to P deficiency, due to biochemical changes, formation of cluster roots and substantial release of carboxylates (Tiziani et al., 2020; Vance et al., 2003). Its high carboxylate exudation also makes it an ideal candidate for studying the role of root exudates in bioweathering processes. However, despite extensive research on exudates and nutrient mobilization, the role of soil development stage and mineralogical context in shaping these interactions remains largely unexplored. In this study, we address this gap by growing *Lupinus albus* under a gradient of P availability, collecting its exudates, and applying these exudates cocktails in dissolution experiments with three contrasting soil horizons representing incipient, intermediate and advanced stages of soil genesis. This design allows us to test how plant responses to nutrient limitation translate into changes in exudate profiles, and how these exudates, in turn, drive bioweathering processes across soils of contrasting mineralogy and weathering degree. In doing so, we link plant-level physiological responses with pedogenesis, providing new insights into how rhizosphere processes may shape nutrient cycling throughout soil development.

## Material and Methods

### Phosphorus gradient experiment

#### Plant cultivation

White lupin (*Lupinus albus*) was grown hydroponically in a growth chamber in October–November 2023. Seeds were surface-sterilized in 30% H□O□ for 10 min (Shane et al., 2003) and germinated in the dark at 25□°C for five days in 1 mM CaCl□ and 5□µM H□BO□ (O’Sullivan et al., 2021). After germination, 100 seedlings were transferred to five 25-L containers with quarter-strength Hoagland solution (Hoagland & Arnon, 1950) for two days, then half-strength Hoagland solution until two fully expanded leaflets developed. Seedlings were then moved to aerated modified Hoagland solutions (**Supplementary material 1**) with KH□PO□ at 5, 10, 20, 30, or 50□µM to generate a P response curve (Abdolzadeh et al., 2010; Keerthisinghe et al., 1998; O’Sullivan et al., 2021). Potassium was balanced with KCl (Pang et al., 2010). Nutrient solutions were renewed every five days, and pH maintained at 6.0. Cotyledons were removed at developmental stage 1.2 (two leaves emerged; J. Walker et al., 2011) to induce early P deficiency.

Plants were grown under a 16-h light/8-h dark cycle at 24□°C/18□°C, 50% RH, and 400□µmol□m□^2^□s□^1^ light intensity. After seven weeks (49 days, floral stage), root exudates were collected, and shoots and roots harvested, dried at 60□°C for 24 h, and weighed. Leaf area was measured with a LI-3000A leaf area meter. Roots from three replicates per P treatment were scanned in a transparent water tray using a flatbed scanner (Epson V850 Pro). Root tips and total root length were analyzed with RhizoVision (Seethepalli & York, 2020), and cluster roots manually delineated using SmartRoots (Lobet et al., 2011).

### Collection and analysis of root exudates

We collected root exudates over two consecutive days. After 4 h of light, roots were washed three times in distilled water, then immersed in 130 mL distilled water in foil-wrapped 150 mL beakers. After 2 h, plants returned to hydroponics. Solutions were stored at 4□°C overnight. Twenty-four hours later, we repeated the process, yielding four 60-mL vials, which were pooled, filtered (0.45□µm), and split into four sub-samples. pH, non-purgeable organic carbon (NPOC), and total nitrogen (TN) were measured using a TOC-L analyzer. Samples were lyophilized; one subsample was used for metabolomics, three for batch dissolution.

For metabolomics, lyophilized material was reconstituted in 2 mL nanopure water, 1 mL analyzed via GC-MS, and 1 mL reserved. Samples were dried in a speed vacuum, derivatized, and analyzed following Xu et al., 2018. Metabolites were identified using an expanded Agilent Fiehn database, NIST23, and Wiley 11th Edition. Absolute quantification was achieved using serial dilutions (0, 0.1, 1, 10, and 100 µg/mL) of standard compounds (aspartic acid, citric acid, fructose, glucose, glutamic acid, malic acid, malonic acid, oxalic acid, succinic acid, sucrose) analyzed in triplicate. Regression values (R^2^) were consistently >0.95.

### Batch dissolution test

#### Soils

We used three soil horizons (“Ae”, “Bf” and “BC”-**Supplementary material 2**) representing different stages of pedogenesis within a podzolic chronosequence near Cox Bay, on the west coast of Vancouver Island, British Columbia (latitude 49° 6′N, longitude 125° 52′W). This chronosequence is developing from beach sand deposits advancing towards the ocean at a rate of 0.26 m per year, providing soils of increasing age with distance from the present shoreline (Singleton & Lavkulich, 1987). Mean annual precipitation is 3,200 mm and the average temperature is 8.9 °C.

A chronosequence is a “space-for-time substitution”, where soils of different ages but similar forming factors (parent material, climate, topography, and organisms (Jenny, 1941)) can be compared to infer the rate of soil development and ecosystem change (Walker et al., 2010) . The soils of the chronosequence ranged from an Orthic Dystric Brunisol (10 meters behind the present beach) to a Humo-Ferric Podzol (100 meters from the present beach) (Vermeire et al., 2016). From these soils, we selected three horizons. *BC* soil horizon was sampled 5 meters behind the present beach and represents the incipient of soil formation, very similar to the parent material, enriched in primary minerals. *Ae*, the eluvial grey horizon and *Bf*, the illuvial brown-red horizon, were sampled from the Humo-Ferric Podzol, 150 meters away from the present beach, respectively at 0–10 cm and 10–30 cm. By selecting these horizons, we ensured that the differences among them reflected pedogenic processes rather than other soil forming factors. Soils were air-dried and sieved to < 2 mm. Three samples of each soil horizon were used to determine major soil characteristics.

#### General soil description

Soil pH, CEC, exchangeable cations, total C and N, col and hot C extraction, available NH_4_^+^ and NO_3_^-^, organic and inorganic P, extractable Fe, Al, Si, Mn, and total elemental concentration were measured using standards protocols (Carter & Gregorich, 2007; Ghani et al., 2003; U.S. Environmental Protection Agency, 1996; T. W. Walker & Adams, 1958; Ziadi & Sen Tran, 2007). Soil mineralogy was assessed by powder X-ray diffraction (XRD) on a Phillips X’Pert Multipurpose X-ray diffractometer. Details protocol available in **Supplementary material 3**. The main properties of the three soil horizons are shown in **Table 1** and in **Supplementary material 4**.

**Table 1:**
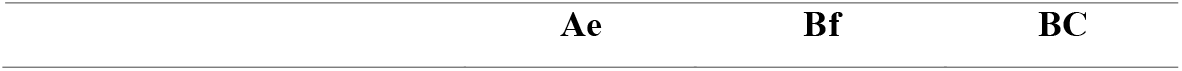
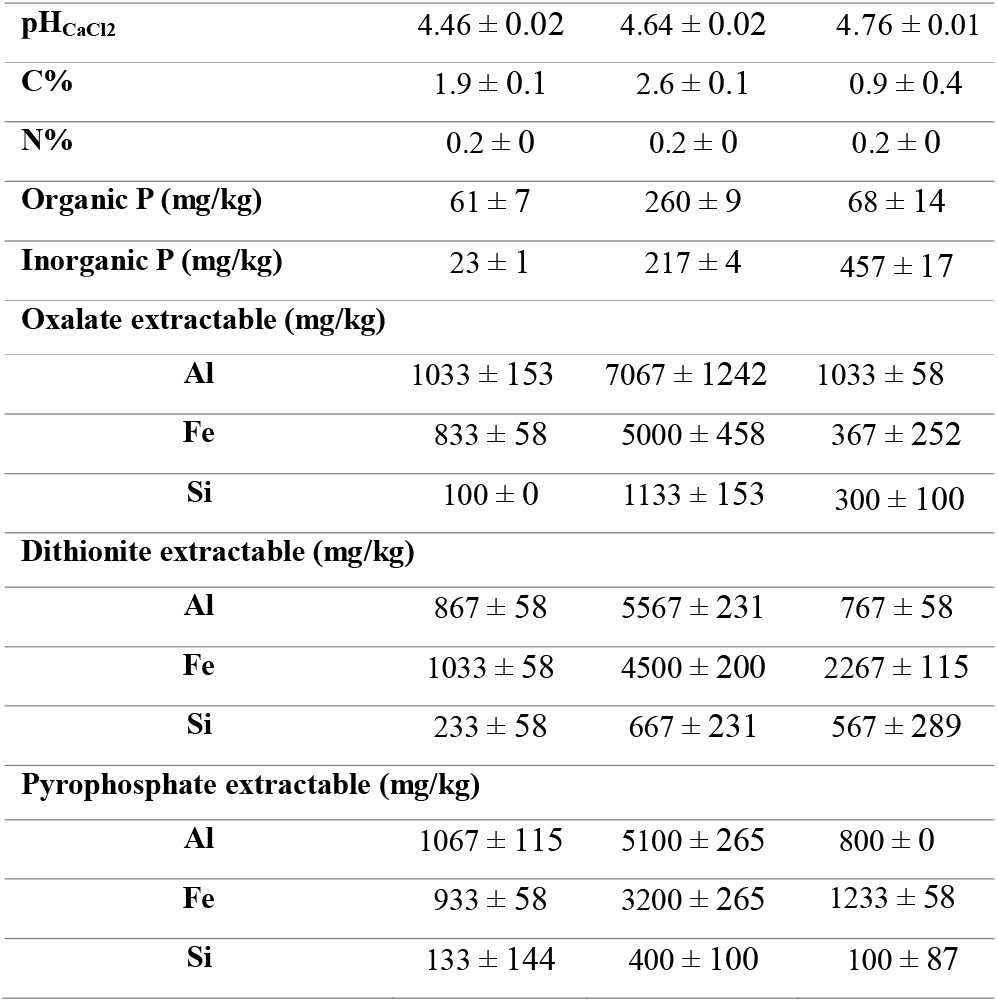
Soil physico-chemical characteristics. Values are presented as mean ± standard deviation.

### Batch dissolution procedure and analysis

Two hundred fifty mg of each soil horizon were weighed in 15 mL Falcon tubes. Sixty mL of the root exudates collected from the plants were freeze-dried, resuspended in 5 mL of MiliQ water, added to the soil and shaken at 100 rpm at room temperature for 24 hours. Solutions were centrifuged (2,500 g, 20 min), filtered (0.45 µm), acidified with 10% HNO□, and analyzed for elemental composition (Mg, Al, Si, P, K, Ca, Fe) by Inductively Coupled Plasma Mass Spectrometry (ICP-MS), and for NPOC/TN by a TOC-L analyzer.

#### Statistical analysis

One-way analysis of variance (ANOVA) assessed P treatment effects on plant traits and root exudate (R Core Team, 2022). Residual normality was tested with Shapiro–Wilk and Q–Q plots and Levene’s test checked variances homogeneity. Tukey’s HSD post hoc test was applied to identify significant differences between group means. Log or square-root transformations were applied if assumptions failed. If assumptions could not be met despite transformation, Welch’s ANOVA and the Games–Howell post hoc test (via the rstatix package (Kassambara A, 2023)) were used instead. The total exudation rate of OA (µg mL^-1^ h^-1^) was computed by summing the exudation rate of the 7 quantified OAs (malic, citric, oxalic, tartaric, succinic, malonic, fumaric acid). Linear regressions determined plant trait contribution to total OA rate. Metabolomics data was analyzed using Partial Least Squares Discriminant Analysis (PLS-DA) from the mixOmics package (Rohart et al., 2017) after log-transformation and mean-centering. Variable importance in projection (VIP) scores greater than 1 identified the most influential metabolites in differentiating treatment groups. Net percentage of element dissolved was calculated as the ratio of element released in the batch dissolution to its total soil concentration (measured by total digestion). Because OA concentrations varied widely across P treatments, root exudates were grouped into three categories (low, intermediate, high) based on the total concentration of the seven OAs, considering the 5 mL volume used to re-suspend lyophilized samples. Total OA concentration, rather than individual OAs, was used to assess overall dissolution power of the exudates. A two-way ANOVA tested the effects of soil horizon (Ae, Bf, BC), OA category (control, low, medium, high), and their interaction on element dissolution. Linear regressions examined relationships between total OA concentration and net dissolved element concentrations, with significance assessed by p-values and R^2^. Additional regressions between selected elements (Al vs. Fe, Si vs. Fe, Ca vs. Mg) were used to explore potential co-dissolution processes from Fe-, Al-oxyhydroxides and Ca-, Mg-silicates within each soil horizon.

## Results

### Plant response to a gradient of phosphorus limitation

#### Plant growth parameters after 7 weeks of growth

Phosphorus supply had a clear impact on white lupin growth and root development after 7 weeks (**Fig. 1, Table 2**). Shoot biomass increased with P concentration and plateaued at intermediate to high P levels (20 to 50 µM P), while root biomass peaked at 20 µM P. As a result, the root:shoot ratio (R:S) decreased significantly with increasing P (**Table 2**). Interestingly, P supply had no significant effect on total root length or the number of root tips.

**Table 2:** Plant parameters of white lupin after 7 weeks of growth under different P treatments. Values are presented as mean ± standard error. Different letters indicate statistically significant differen on Tukey’s HSD test (p < 0.05).

|  | Shoot biomass |  |  |  | Root biomass |  |  |  | Root: Shoot (RS) |  |  |  | Total root length |  |  |  | Total length of cluster roots |  |  |  | Number of cluster roots |  |  |  | Cluster root length normalized by total root length |  |  |  | Number of root tips |  |  |  |
| --- | --- | --- | --- | --- | --- | --- | --- | --- | --- | --- | --- | --- | --- | --- | --- | --- | --- | --- | --- | --- | --- | --- | --- | --- | --- | --- | --- | --- | --- | --- | --- | --- |
|  | (mg) |  |  |  | (mg) |  |  |  | (-) |  |  |  | (mm) |  |  |  | (mm) |  |  |  |  |  |  |  | % |  |  |  |  |  |  |  |
| $\mu\text{M P}$ | 507.4 | ± | 59.2 | c | 321.8 | ± | 42.9 | b | 0.63 | ± | 0.04 | a | 748.2 | ± | 65.8 | a | 81.0 | ± | 7.1 | a | 57 | ± | 4 | a | 10.8 | ± | 0.1 | a | 396.0 | ± | 38.5 | a |
| $\mu\text{M P}$ | 701.9 | ± | 55.2 | bc | 519.2 | ± | 24.4 | ab | 0.75 | ± | 0.05 | a | 982.5 | ± | 112.6 | a | 96.0 | ± | 11.2 | a | 75 | ± | 8 | a | 9.9 | ± | 0.7 | a | 541.2 | ± | 92.3 | a |
| $\mu\text{M P}$ | 1517.9 | ± | 192.9 | ab | 868.3 | ± | 165.7 | a | 0.56 | ± | 0.04 | ab | 1438.9 | ± | 127.9 | a | 98.2 | ± | 9.6 | a | 77 | ± | 7 | a | 7.0 | ± | 0.9 | ab | 559.3 | ± | 39.3 | a |
| $\mu\text{M P}$ | 1636.1 | ± | 327.7 | a | 615.2 | ± | 120.9 | ab | 0.38 | ± | 0.03 | bc | 1140.7 | ± | 240.9 | a | 81.8 | ± | 25.6 | a | 67 | ± | 16 | a | 7.2 | ± | 1.2 | ab | 425.7 | ± | 68.9 | a |
| $\mu\text{M P}$ | 1156.2 | ± | 232.7 | ab | 411.4 | ± | 114.7 | ab | 0.34 | ± | 0.04 | c | 778.2 | ± | 275.7 | a | 19.2 | ± | 3.4 | b | 15 | ± | 3 | b | 3.3 | ± | 0.8 | b | 242.5 | ± | 65.2 | a |

**Figure 1.**
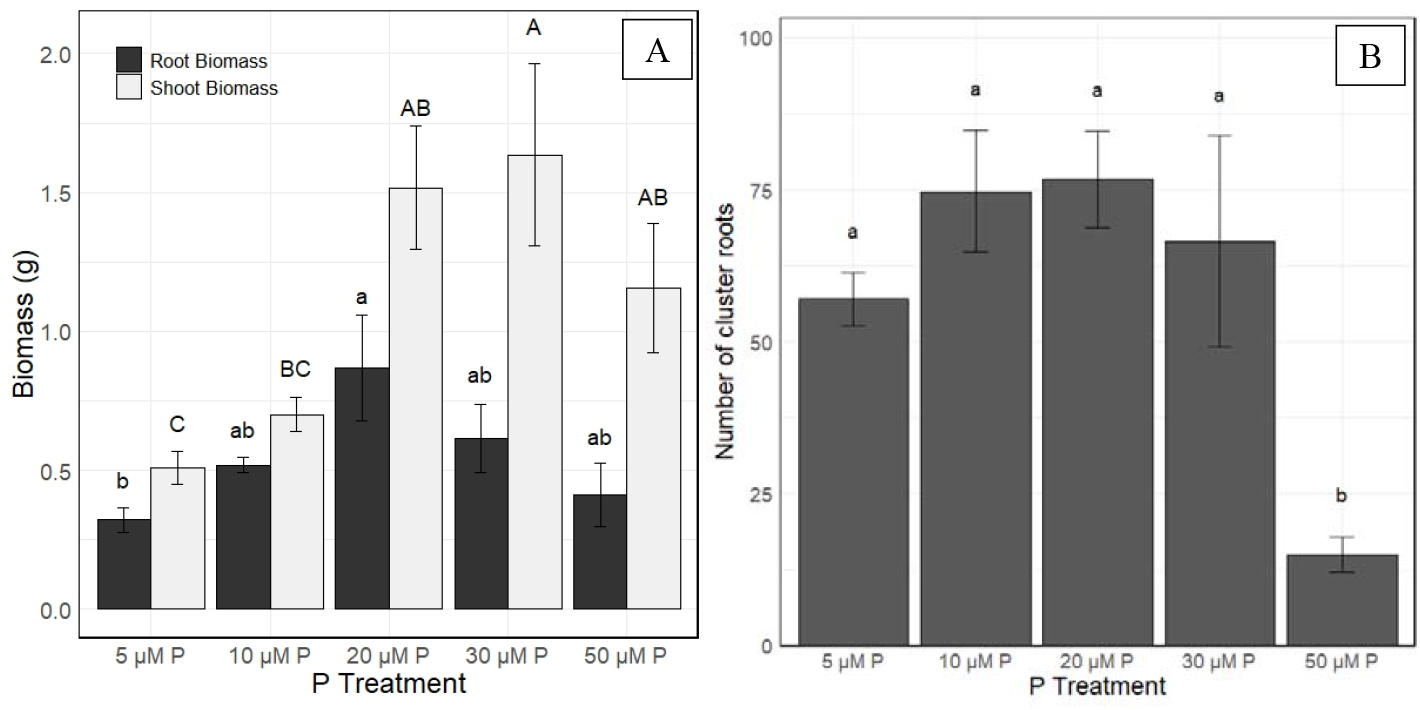
A. Shoot and root biomass (g). Different lowercase letters denote significant differences in root biomass among the P treatments. Different uppercase letter denotes significant differences in shoot biomass. B Number of cluster roots. Bars represent mean ± SE.

Cluster roots were observed in all P treatments, but their development was strongly reduced at high P (50 µM), both in number and length (**Fig. 1B, Fig. 2**). The proportion of cluster roots relative to total root length was significantly lower at 50 µM P, compared to the low P treatments (5 and 10 µM).

**Figure 2.**
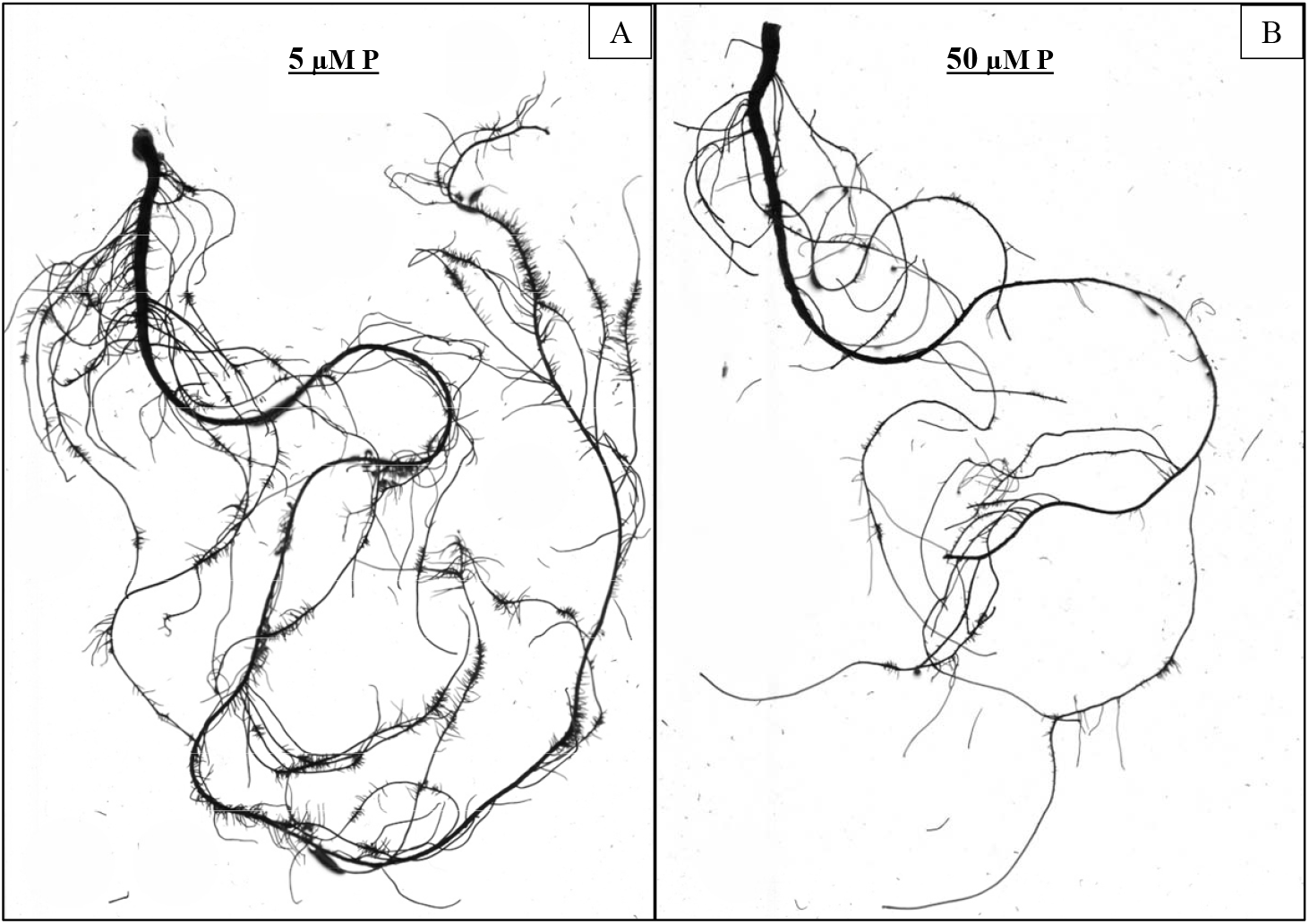
Representative root systems of white lupin after 7 weeks of growth under two P treatments: (A) 5 µM P (lowest P level) and (B) 50 µM P (highest P level).

#### White lupin root exudation in a phosphorus gradient

Root exudate pH, NPOC and total OA showed clear trends across P treatments (**Table 3**). The pH of the root exudate solution peaked at 30 µM P (5.69), significantly higher that at 10 µM P (4.92), with a positive linear relationship with root biomass (R^2^ = 0.495). NPOC followed a similar Gaussian trend, highest at 30 µM P (6.83 mg/L) and lowest at 5 µM P (3.37 mg/L), also positively correlated with root biomass (R^2^ = 0.555). When normalized by root biomass, NPOC levels were highly variable and showed no clear pattern.

**Table 3:** Characteristics of root exudate solutions from white lupin grown under different P treatments. Values are presented as mean ± standard error. Different letters indicate statistically differences among treatments based on Tukey’s HSD test (p < 0.05). NPOC = non-purgeable organic carbon, OA = organic acid, TN= total nitrogen.

|  | pH |  |  |  | NPOC |  |  |  | NPOC per g root biomass |  |  |  | Total OA exudation rate |  |  |  | Total OA exudation rate per g root biomass |  |  |  | TN |  |  |  |
| --- | --- | --- | --- | --- | --- | --- | --- | --- | --- | --- | --- | --- | --- | --- | --- | --- | --- | --- | --- | --- | --- | --- | --- | --- |
|  |  |  |  |  | (mg L<sup>-1</sup>h<sup>-1</sup>) |  |  |  | (mg L<sup>-1</sup>h<sup>-1</sup> g root<sup>-1</sup>) |  |  |  | (μg L<sup>-1</sup>h<sup>-1</sup>) |  |  |  | (μg L<sup>-1</sup>h<sup>-1</sup> g root<sup>-1</sup>) |  |  |  | (mg L<sup>-1</sup>h<sup>-1</sup>) |  |  |  |
| 05 μM P | 3.37 | ± | 0.35 | ab | 1.68 | ± | 0.17 | b | 5.81 | ± | 1.13 | a | 0.0037 | ± | 0.00062 | b | 0.0123 | ± | 0.00238 | a | 0.73 | ± | 0.05 | a |
| 10 μM P | 3.72 | ± | 0.55 | b | 1.86 | ± | 0.28 | b | 4.61 | ± | 0.90 | a | 0.0036 | ± | 0.00059 | b | 0.00701 | ± | 0.00105 | a | 0.66 | ± | 0.16 | a |
| 20 μM P | 5.96 | ± | 0.37 | ab | 2.98 | ± | 0.19 | ab | 4.71 | ± | 0.26 | a | 0.0059 | ± | 0.00115 | ab | 0.00721 | ± | 0.00193 | a | 1.08 | ± | 0.52 | a |
| 30 μM P | 6.83 | ± | 1.14 | a | 3.41 | ± | 0.57 | a | 5.76 | ± | 0.39 | a | 0.0068 | ± | 0.00129 | a | 0.0119 | ± | 0.00205 | a | 1.15 | ± | 0.38 | a |
| 50 μM P | 3.43 | ± | 0.23 | ab | 1.71 | ± | 0.12 | b | 5.23 | ± | 1.28 | a | 0.0035 | ± | 0.00107 | b | 0.00878 | ± | 0.00125 | a | 0.48 | ± | 0.08 | a |

Total exudation rate of OA followed the same bell-shaped trend, highest at 30 µM P (0.0068 μg mL□^1^ h□^1^) and lowest at 50 µM P (0.0035 μg L^-1^h^-1^), with marginal differences among treatments (*p* = 0.0707). Normalized rates were not significantly different but tended to be higher at 5 and 30 µM P. Total OA exudation was positively correlated with leaf biomass, leaf area, and root biomass (R^2^ = 0.534, 0.500, and 0.421, respectively, **Supplementary Material 5**)

Oxalic acid was the dominant measured OA exuded at 7 weeks, ranging from 0.00123 µg mL□^1^ h□^1^ (10 µM P) to 0.00272 µg mL□^1^ h□^1^ (30 µM P), and accounting for up to 42.3% of the total OA pool (at 20 µM P). Succinic acid was the second most abundant, particularly at higher P (up to 23.2% at 50 µM P). While oxalic, succinic, and malonic acid levels were relatively stable across treatments, other OAs varied significantly with P availability (**Fig. 3**). Citric and fumaric acid exudation peaked at 30 µM P compared to 10 µM P, while malic acid was highest at 30 µM P, significantly exceeding levels at 5, 10, and 50 µM P. Tartaric acid was detected in all treatments except 50 µM P. Sugars (glucose, sucrose, fructose) and amino acids (L-glutamic acid, L-aspartic acid) showed no significant differences among P treatments.

**Figure 3.**
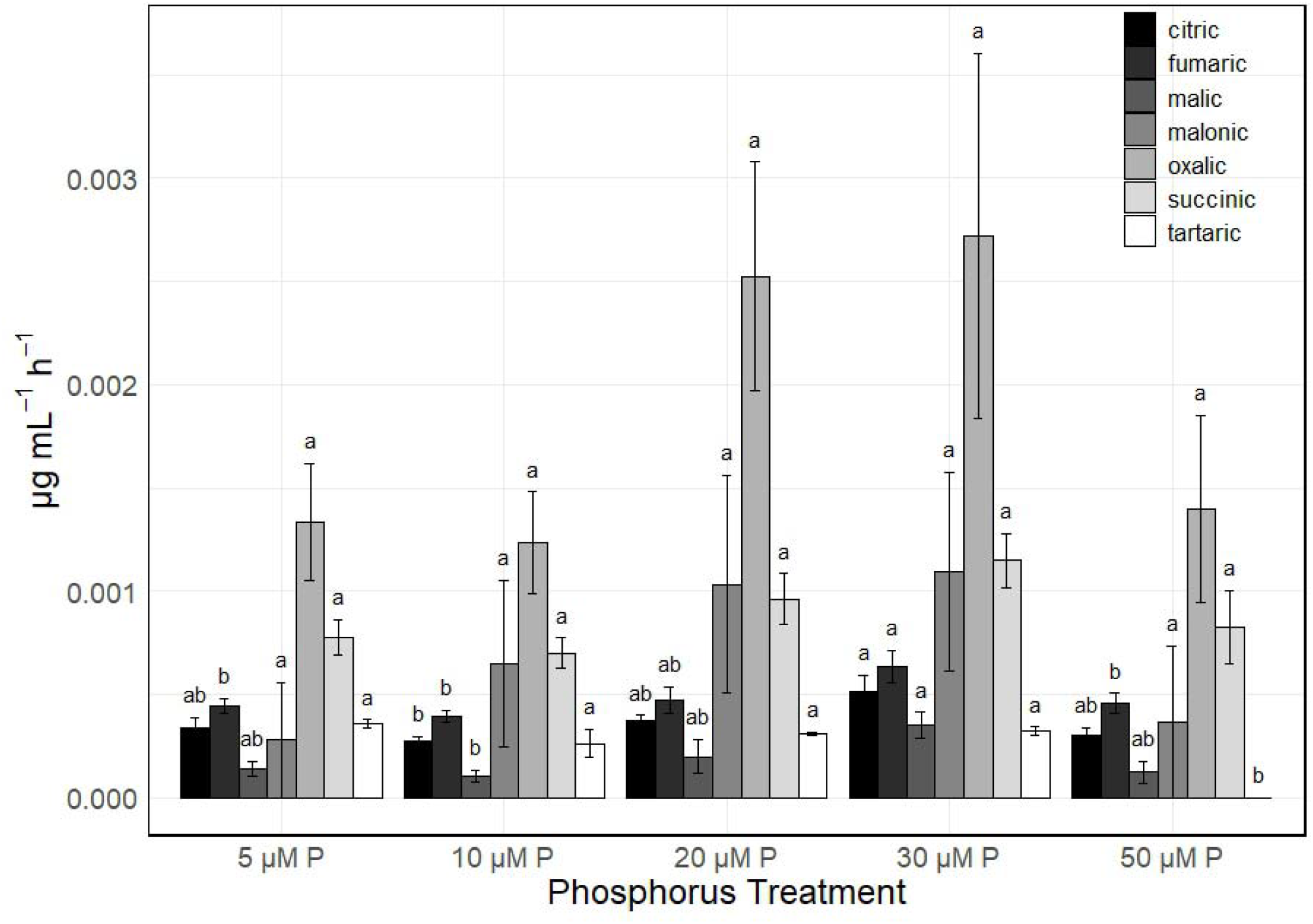
Organic acid concentrations in root exudates of white lupin grown under different P treatments. Bars represent mean ± standard error. Different letters within each organic acid group indicate statistically significant differences among P treatments (Tukey’s HSD test, p < 0.05).

A total of 182 metabolites were detected in the root exudates, including 103 identified compounds and 79 unknowns. Partial Least Squares Discriminant Analysis (PLS-DA) revealed a better separation between two low P treatments (05 µM P and 10 µM P), and the 3 higher P treatments (20, 30 and 50 µM P) which clustered closely (**Fig. 4**). The first two PLS-DA components explained 30% and 15% of the total variance, respectively. In total, 71 metabolites differed significantly among the P treatments (p < 0.05), comprising 45 identified and 26 unknown compounds.

**Figure 4.**
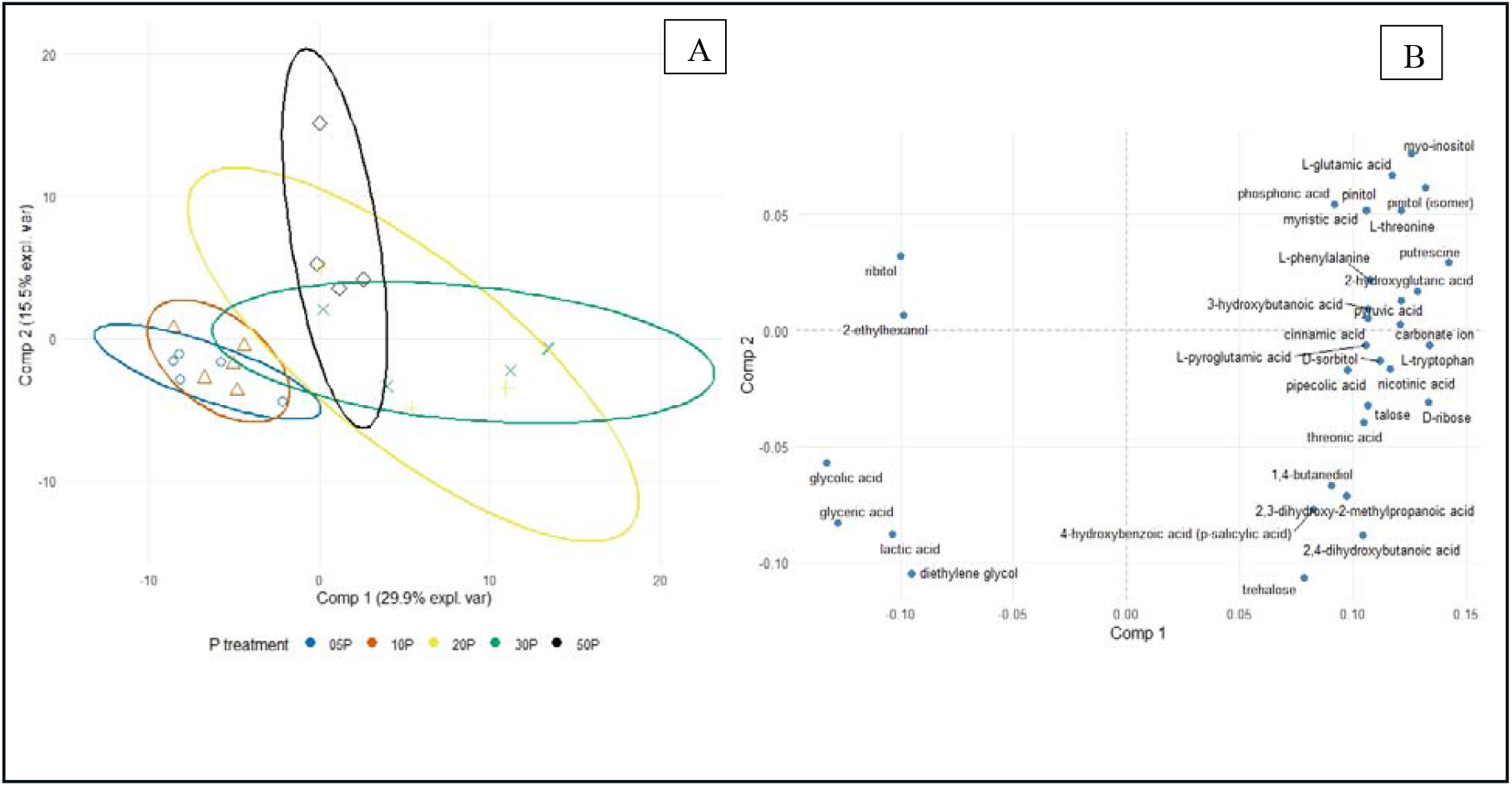
(A) PLS-DA score plot showing the distribution of white lupin exudate samples based on metabolite profiles measured by GC-MS. Ellipses sent 95% confidence intervals for each P treatment group. Component 1 explains 29.9% of the variance, and Component 2 explains 15.5%. orrelation circle plot showing the most influential metabolites (VIP > 1) contributing to group separation.

### Batch dissolution test

#### Mineralogical properties of the soil horizons

The BC horizon was rich in primary minerals such as hornblende and chlorite (**Fig. 5**), similar to the beach parent material. It contained intermediate levels of crystalline Fe oxides (Fe_dcb_) and low levels of crystalline Al (Al_dcb_) and exchangeable Al (**Table 1**). The differences between Al_ox_ and Al_pyr_, and between Fe_ox_ and Fe_pyr_, indicated the presence of amorphous forms of Al and Fe, with a higher proportion of amorphous Fe.

**Figure 5.**
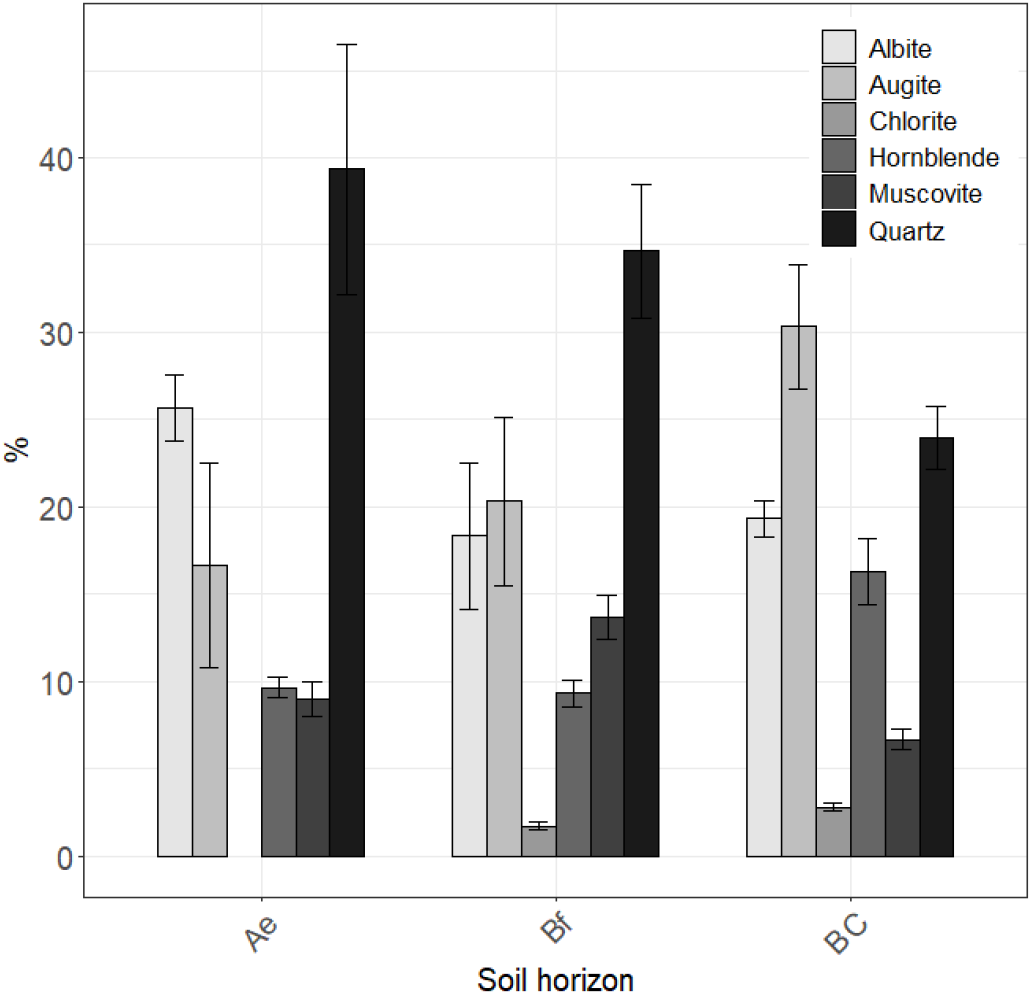
Mineralogical composition of the three soil horizons. Bars represent mean ± standard error.

The Ae horizon showed enrichment in quartz and albite, and the presence of secondary clay minerals. Chlorite and serpentine were absent. Low levels of Al_dcb_, Fe_dcb_, and Si_dcb_ suggested depletion of Fe, Al, and Si oxides. The Fe_ox_:Fe_dcb_ ratio below 1, indicated a greater proportion of crystalline Fe forms relative to amorphous.

The Bf horizon displayed intermediate mineralogical characteristics to the BC and Ae soil horizons. It was characterized by high Al_dcb_, Fe_dcb_, and Si_dcb_ values. Oxalate extractions showed high amounts of amorphous and organically complexed Al, Fe, and Si. A large difference between Al_ox_ and Al_pyr_, along with a high Al_ox_:Al_dcb_ ratio (∼6), reflects a predominance of amorphous Al.

#### Root exudates impact on mineral dissolution

Addition of root exudates significantly increased the release of Mg, Al, Si, P, K, Ca, and Fe compared with the water control (**Fig. 6; Supplementary Material 6**). For Al and Fe, effects varied with soil horizon: negligible in Ae, limited to high OA concentrations in BC, but strong across all concentrations in Bf (**Fig. 7**). Medium and high OA concentrations also enhanced Si, K, and P release relative to low OA in all horizons. Total OA concentration correlated significantly with the dissolution of K (R^2^ = 0.28), P (R^2^ = 0.26), Si (R^2^ = 0.11), and Fe (R^2^ = 0.08), but not with Ca or Mg.

**Figure 6.**
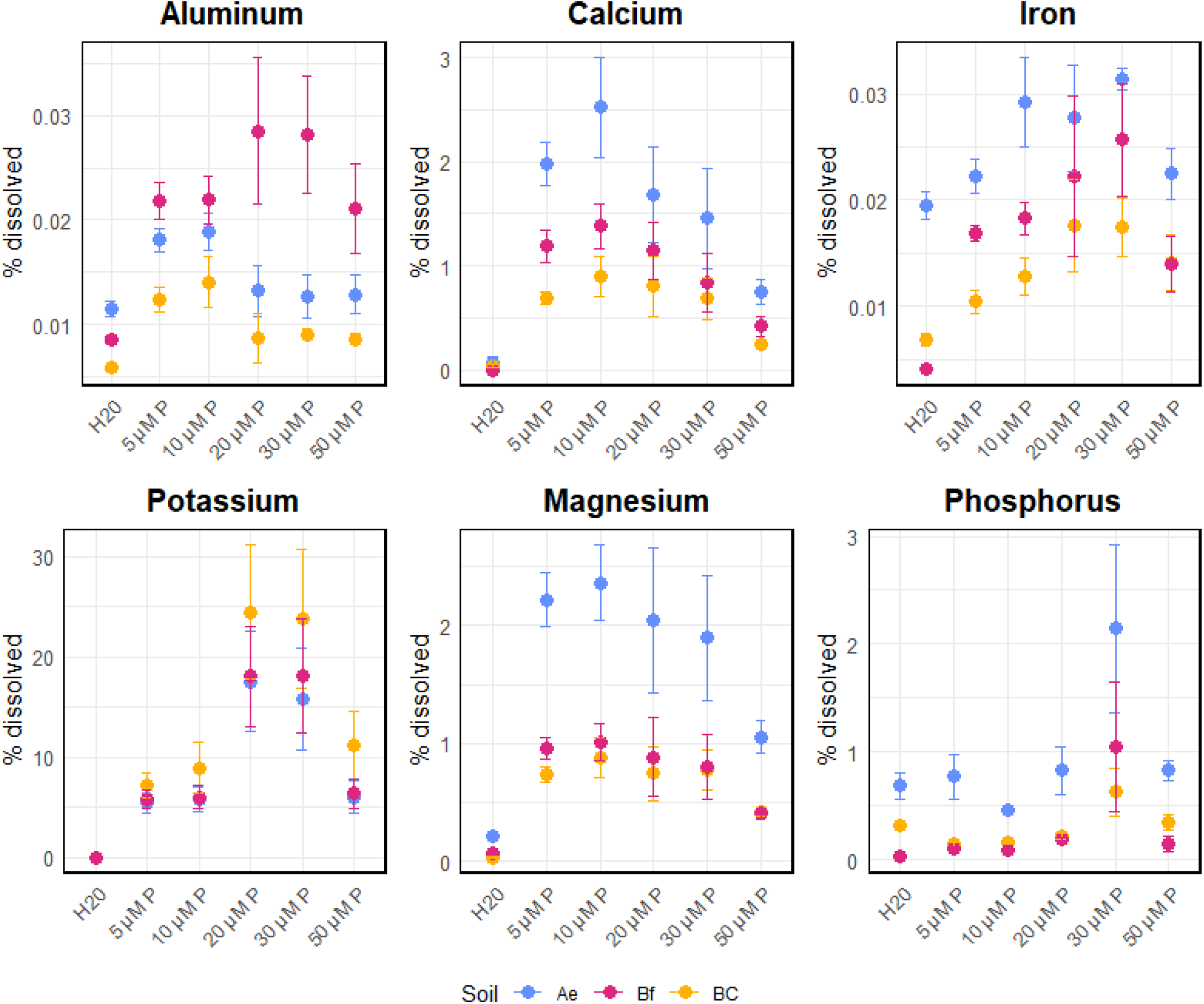
Net percentage of each element dissolved, calculated as the ratio between the concentration of the element released during the batch dissolution experiment and its total concentration in soil (measured by total digestion). Results are grouped by P treatment, with H□ O serving as the control. Data points are color-coded by soil horizon. Values are presented as means ± standard error.

**Figure 7.**
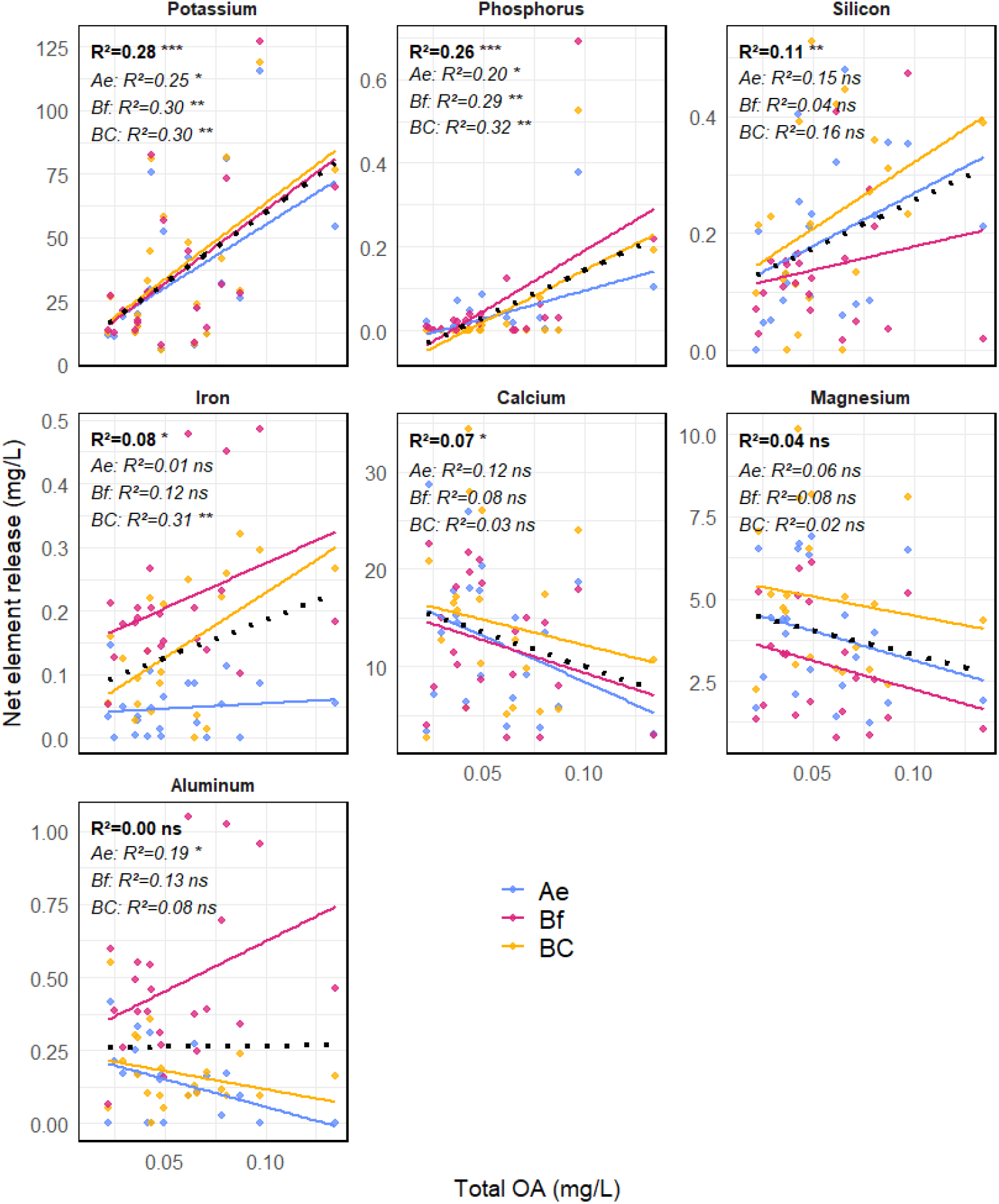
Linear relationships between the net element release and the total organic acid concentration in white lupin exudates for seven elements measured in this study. Data points are color-coded by soil type. Linear regressions were fitted both across the entire dataset (dotted regression line) and separately for each soil horizon (regression lines of different colors). The displayed R^2^ and significance (p<0.05 *; p<0.01**; p < 0.001***, p>0.05 ns) in bold correspond to the regressions fitted across all data points whereas italic display the R^2^ and significance for each soil type.

**Figure 8.**
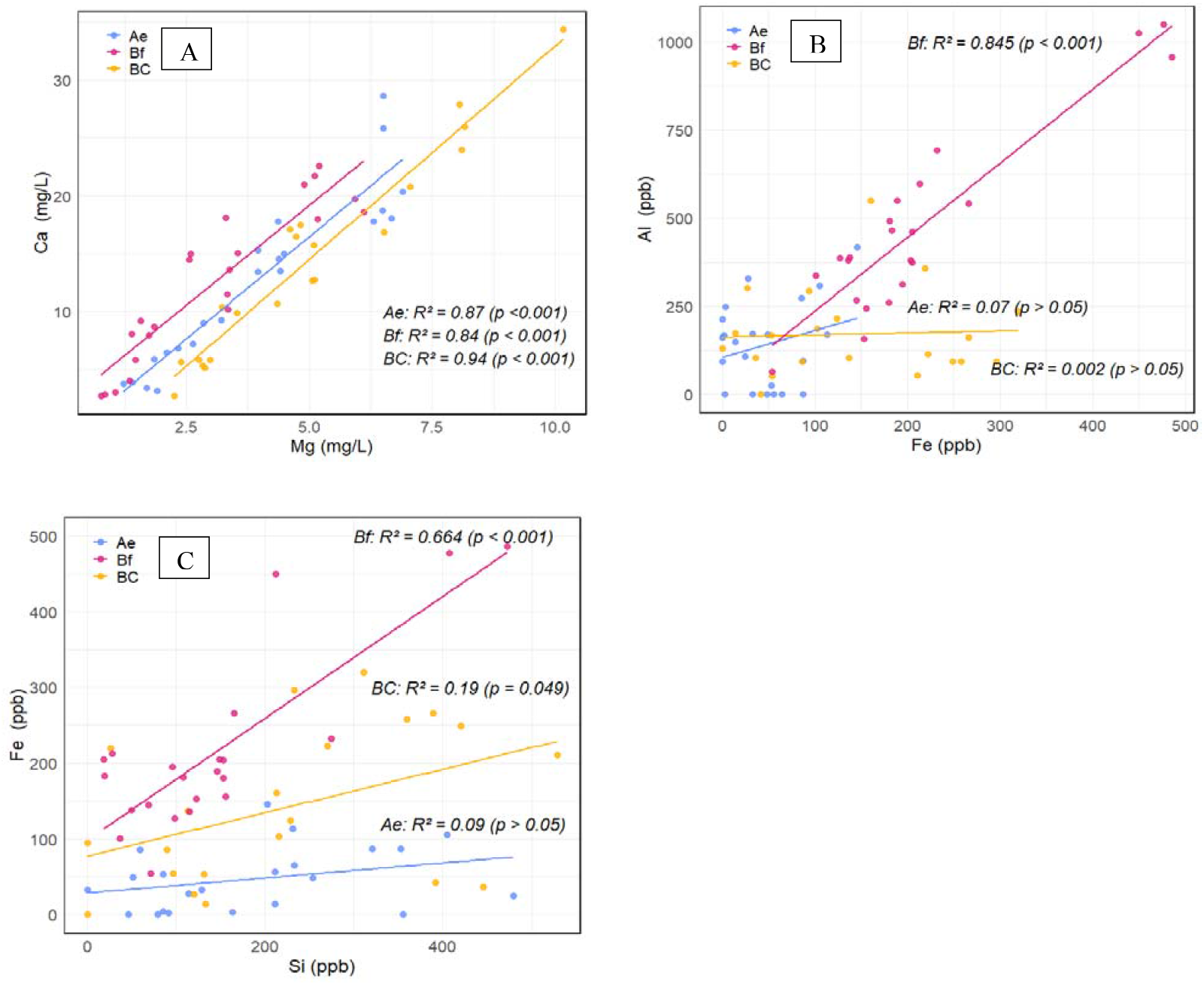
Co-solubilization process between elements in the three soil horizons. A) Mg-Ca, B) Fe-Al, and C) Si-Fe. Data points are color-coded according to the three different soil types. Linear regression between the elements were performed separately for each soil horizon. R^2^ and p-values (p) indicate the strength and significance of the regressions.

Element co-dissolution patterns further highlighted horizon-specific dynamics (**Fig. 8**). Fe and Al release were strongly correlated in the Bf horizon, but not in Ae or BC. Si–Fe relationships followed the same trend— strongest in Bf, moderate in BC, and weak in Ae. Ca and Mg release showed consistent positive correlations across all horizons. Regression details (slopes, intercepts, p-values) are reported in **Supplementary Material 7**.

Net element dissolution was shaped by both P treatment and soil horizon (**Fig. 6**). Most elements showed a Gaussian response, with lowest dissolution at 5 and 50 µM P and highest at 20–30 µM. Release was greatest in the Ae horizon and lowest in BC, while Bf was intermediate except for Al, which dissolved most strongly there.

#### How does P gradient influence plant parameters and root exudates?

Our results reveal a clear tipping point in plant response along the P gradient. Despite some variability in biomass data, P concentrations of 20–30 µM P supported shoot biomass comparable to 50 µM P, the level reported as optimal for white lupin. In this range, however, root biomass was higher, and root-to-shoot ratios increased, suggesting enhanced C transfer to the roots—a well-documented response to nutrient limitation (Hermans et al., 2006). At 50 µM P, cluster root formation was significantly reduced, confirming sufficient P supply. Thus, mild P limitation (30 µM, ∼60% of recommended dose) appears to trigger adaptive root traits without reducing shoot growth, consistent with of Keerthisinghe et al., 1998, who noted that cluster root formation does not necessarily compromise biomass production (C. Wang et al., 2025).

Another well-established response of white lupin to P limitation is increased OA exudation. Of the seven OAs quantified, only citric, fumaric, malic, and tartaric acids varied significantly across treatments. Both NPOC and total OA peaked at 30 µM P, suggesting that moderate P limitation enhances exudation without affecting shoot biomass. When expressed per gram of dry root weight, total OA and NPOC levels—despite high variability— tended to be highest at 5 µM P and 30 µM P, but likely for different reasons: greater root biomass at 30 µM P versus higher exudation per unit root at 5 µM P.

Similar to the findings of Mimmo et al., 2011, oxalic acid was the dominant compound in our samples, even though most other studies typically report citric and malic acids as the major carboxylates exuded by white lupin (Funayama-Noguchi et al., 2015; Neumann & Römheld, 1999; Wang et al., 2013). This difference may be due to the timing of our sampling: root exudates were collected 49 days after emergence, during the flowering stage. Previous work has shown that exudation dynamics are closely linked to plant developmental stages. For example, O’Sullivan et al., 2022 reported that exudation rates tend to peak during early growth but decline during reproductive stages as photosynthates are increasingly allocated to biomass accumulation and grain development (Haase et al., 2007). Similarly, Shen et al., 2003 observed a cyclic pattern in citrate exudation under P-limited conditions, with the highest levels occurring before flowering. Together, these findings suggest that exudation quantity and composition are shaped by both P availability and plant developmental stage.

In this study, we applied metabolomics to profile root exudates of white lupin for the first time. Higher variability was observed among replicates under medium-to-high P (20–50 µM) compared to low P (5–10 µM). Multivariate analyses revealed two distinct clusters separating low P from higher P treatments. Beyond carboxylates, several metabolites showed treatment-specific patterns, including myo-inositol, a known exudate component. These results highlight a release of a broader array of metabolites in white lupin exudates in response to changes in P availability—beyond the well-known carboxylate response. This is well aligned with the findings from Müller et al., 2015 that highlight a clear metabolomics shift in white lupin P-deficient shoots and roots. Overall, our metabolomic and physiological findings support a threshold around 20 µM P that distinguishes low P conditions from intermediate-to-high supply, associated with a shift in both root traits and exudation profiles.

#### How does organic acid exudation influence bioweathering processes?

To our knowledge, this is the first study to use realistic exudate cocktails and OA concentration to investigate bioweathering processes. Previous batch dissolution experiments typically applied synthetic or isolated compounds to soils or minerals (Garcia Arredondo et al., 2019; Jones, 1998; Oburger et al., 2009, 2011), often at relatively high ligand concentration (∼0.5 mM) and solid:liquid ratios of 2.5-100 g/L (Edayilam et al., 2018; Jones & Darrah, 1994; D. Wang et al., 2016). Here, we used 50 g/L (1:20), closer to soil conditions, but our exudate solutions contained much lower OA levels: maximum malic and oxalic acid concentrations were 0.00058 and 0.0734 mg/L, respectively—over 100-fold lower than typical experimental levels (e.g., 67.05 mg/L malic acid, 45.015 mg/L oxalic acid at 0.5 mM). Thus, this study highlights bioweathering processes occurring at OA concentrations far below those usually tested.

Our metabolomic analysis revealed more than 100 metabolites in lupin exudates, implying that interactive effects likely enhanced dissolution compared with single OA. Indeed, Oburger et al., 2009 showed that citrate, malate, and oxalate mobilized more P in combination with malonate than individually. Consistent with this, we found that total OA concentration was a stronger predictor of elemental release than individual OAs. Given the very low OA levels observed, it is likely that additional metabolites also contributed to metal release.

#### How do bioweathering processes vary depending on the soil physicochemical context?

The soil horizons used in batch dissolution experiments represent different stages of soil genesis with distinct physicochemical characteristics. White lupin root exudates consistently enhanced the release of Al, Ca, Fe, K, Mg, P, and Si compared with water-only controls, though the magnitude varied by horizon and element, consistent with the specificity of rhizosphere-mediated mobilization (Jones, 1998; Jones & Darrah, 1994). Across all horizons, total OA concentration correlated positively with K, P, and Si release, and with Al and Fe in some cases (BC and Bf), suggesting a strong role for low-molecular-weight OAs in the mobilization of these elements through complexation or ligand-promoted dissolution (Dakora & Phillips, 2002).

No correlation was observed for Ca and Mg, consistent with Oburger et al., 2011, who reported that Ca solubility is mainly pH-driven at 500 µM. In contrast, Jones & Darrah, 1994 found enhanced Ca release with citric or malic acid. In our study, Ca and Mg showed strongly correlated dissolution across horizons, suggesting shared mechanisms, likely from Ca/Mg-bearing silicates (hornblende, augite, chlorite) or organic complexes, as supported by pyrophosphate extraction data. Despite being the most weathered, the Ae horizon released the most Ca and Mg, probably due to its high content of hot-water-extractable C. We hypothesize that the primary source of these elements is organically bound rather than mineral, as fresh OAs mainly remobilize Ca and Mg previously chelated with organic matter in the soil (Rowley et al., 2018).

Potassium release showed a consistent linear relationship with OA concentration across all horizons, suggesting that K mobilization is governed by comparable chemical processes across distinct pedogenetic stages. One potential mechanism for this release is the OA-mediated weathering of K-bearing minerals, as oxalic and citric acids are known to enhance muscovite dissolution (Song & Huang, 1988). Release may also derive from nonexchangeable K (NEK) pools held in mica interlayers (Song & Huang, 1988; Wang et al., 2011; Wang et al., 2000). Both muscovite and hornblende were present in all horizons, and net K release reached ∼20–25% of total mineral K at 20–30 µM P, suggesting that moderate root-derived carboxylates can substantially enhance the mobilization of K, likely through a combination of direct mineral dissolution and desorption from NEK pools.

The effect of OA concentration on Al and Fe release varied across soil horizons, reflecting differences in mineralogical composition and weathering status associated with podzolization. In the Ae horizon, prolonged weathering had depleted reactive Fe and Al oxides (Feox:Fedcb < 1; Blume & Schwertmann, 1969; Rennert et al., 2021) explaining the negligible OA effect. In the BC horizon, representing incipient podzol development, only high concentrations of OAs enhanced Fe release, while no significant effect was observed for Al. Earlier studies suggest that at the initial stages of podzolization, Al tends to be more mobile than Fe due to its greater affinity for forming soluble organo-metallic complexes, whereas Fe precipitates more readily (Jansen et al., 2005; Lundstrom et al., 2000). We hypothesize that this differential behavior in our study could be related to the higher total Fe content compared to Al, as indicated by the different extractable fractions (Fe_dcb_, Fe_pyr_, and Fe_ox_).

In the Bf horizon, both Al and Fe were strongly correlated with the concentration of OAs. The addition of organic C compounds reduced the metal-to-C ratio, enhancing Al solubility from pre-existing organic-metal complexes. An Al:Fe ratio of 2.5 suggests Al-rich amorphous phases, possibly gibbsite-like minerals or non-crystalline Al-organic complexes (Tee Boon et al., 1987), also supported by a high Al_ox_:Al_dcb_ ratio (∼6) and a notable difference between Al_pyr_ and Al_ox_ (Blume & Schwertmann, 1969). These Al-organic complexes are known to be more soluble than their Fe counterparts (Parfitt & Childs, 1988), which may explain the higher responsiveness of Al to increasing OA concentrations.

Al–Si co-solubilization also varied with horizon. In the Bf horizon, OA-driven complexolysis of Fe oxides released Si, likely from short-range-order aluminosilicates (imogolite, proto-allophane) or from Fe- and Al-bound complexes, which are destabilized as organic ligands disrupt metal–oxide bonds (de Tombeur et al., 2021). Similar but weaker patterns occurred in the BC horizon, while no co-release was detected in the Ae horizon due to advanced weathering and depletion of reactive substrates.

Finally, P release was minimal (< 2%) in all horizons, mostly coming from the 30 µM P plants. This likely reflects the very low OA concentrations in our exudate cocktails, far below the thresholds previously shown to mobilize substantial P (1.13 g/L of malic acid and 765 mg/L of oxalic acid; Gerke et al., 2000; Oburger et al., 2009).

### Implications and perspectives

This study aimed to investigate how the intensity of P limitation shapes root exudation and, in turn, bioweathering processes that contribute to nutrient acquisition. Under nutrient-limiting conditions, leaf growth is often more constrained than photosynthesis, resulting in surplus carbohydrates that are redirected to sink organs such as roots, or released into the rhizosphere as root exudates (Prescott et al., 2020). Exudate composition varies with the type of nutrient deficiency (Carvalhais et al., 2011), suggesting that shoot nutrient stoichiometry triggers cascading metabolic and physiological responses that ultimately alter root exudate profiles and uniquely shape rhizosphere dynamics.

Harnessing surplus C and associated biochemical and morphological shifts presents an opportunity to enhance nutrient acquisition in agroecosystems. Specifically, increased root exudation of OAs under P limitation may accelerate bioweathering processes, but responses are soil dependent. First, the intensity of P limitation depends on soil properties, and second, OA effectiveness in promoting mineral weathering varies by soil type. In our podzol soil, nutrient release was largely driven by organic matter dynamics, organo-metallic complexes, and short-range minerals. Although it is often stated that white lupin enhances P acquisition by releasing OAs from cluster roots, our findings showed only modest P mobilization from the soil horizons tested, with a more substantial release of other elements (e.g., K, Fe, Ca). This suggests a need to reinterpret the function of OA exudation—not simply as a targeted strategy for P acquisition, but as part of a broader cascade of biochemical responses, driven by systemic and local cues (e.g., sucrose signaling), rather than solely by the “goal” of P mobilization (Gould & Lewontin, 1979).

Our research shows that applying 60% of the conventional P dose preserved shoot biomass while stimulating belowground responses such as cluster root development and OA exudation at the flowering stage in white lupin. These belowground dynamics reflect whole-plant C flow and source-sink relationships and must be evaluated across plant and soil developmental stages. Overall, plant response to nutrient limitation—and its ecological benefits through bioweathering—depends critically on soil physicochemical properties, which determine both the strength of plant responses and the capacity of exudates to enhance nutrient release.

## Statements & Declarations

We would like to thank colleagues from the University of British Columbia, particularly Fiona Farag and Mary Clare Kendrick for their help in experimental set up. We also thank colleagues from the Environmental Molecular Science Laboratory, specifically Tamas Varga for XRD work, Thomas Wietsma for ICP-MS analysis and Amir Ahkami for their help in the study design. Sasha Pollet is supported by the Wallonia-Brussels International scholarship. This project [GIR003] is supported by the Genomic Innovation for Regenerative Agriculture, Food and Fisheries (GIRAFF) Program, which is funded by the Governments of British Columbia and Canada and delivered by Genome British Columbia and the Investment Agriculture Foundation of BC A portion of this research was performed under the Large-Scale Research program (10.46936/lser.proj.2022.60372/60008596) at the Environmental Molecular Sciences Laboratory located in Pacific Northwest National Laboratory (PNNL), a Department of Energy (DOE) Office of Science User Facility sponsored by the Biological and Environmental Research (BER) program. Battelle operates PNNL for DOE under Contract No. DE-AC05-76RL01830. The authors have no relevant financial or non-financial interests to disclose.

