## Supplementary material for "Phosphorus limitation enhances root exudation and mineral bioweathering across diverse soil process domains"

**Supplementary material 1:** Modified Hoagland recipe

Nutrient solution recipe:

Recipe of full-strength nutrient solution per L of distilled water.

- 1.29 g/L CaNO_3_ NH_4_NO_3_ 10H_2_O
- 5 mL 1M KNO_3_ (Stock solution : 1M = 101.10 g per L)
- 2 mL 1M MgSO_4_ 6H_2_O (Stock solution : 1M = 120.37 g per L).
- 1 mL of micronutrient solution
- Stock solution 1L of micronutrient solution  
  2.85 g H_3_BO_3_ 
  1.81 g MnCl_2_ 4H_2_0 
  0.22 g ZnSO_4_ 7H_2_0 
  0.08 g CuSO4 5H_2_0 
  0.02 g Na_2_Mo0_4_ 2H_2_O 
  3.67 g FeNaEDTA

For the P gradient: add the following solution for 5, 10, 20, 30 and 50 μM P solution per L of distilled water.

5 μM P

- 1 mL KCl (Stock solution: 76.072 g per L)
- 1 mL KH2PO4 (Stock solution: 0.682 g per L)

10 μM P

- 1 mL KCl (Stock solution: 75.68 g per L)
- 1 mL KH2PO4 (Stock solution: 1.364 g per L)

20 μM P

- 1 mL KCl (Stock solution: 74.936 g per L)
- 1 mL KH2PO4 (Stock solution: 2.728 g per L)

30 μM P

- 1 mL KCl (Stock solution: 74.188 g per L)
- 1 mL KH2PO4 (Stock solution: 4.092 g per L)

50 μM P

- 1 mL KCl (Stock solution: 72.688 g per L)
- 1 mL KH2PO4 (Stock solution: 6.82 g per L)

**Supplementary material 2:** Three horizons (BC, Ae, Bh) sampled from the Cox’s Bay chronosequence (from Cornelis et al., 2014)

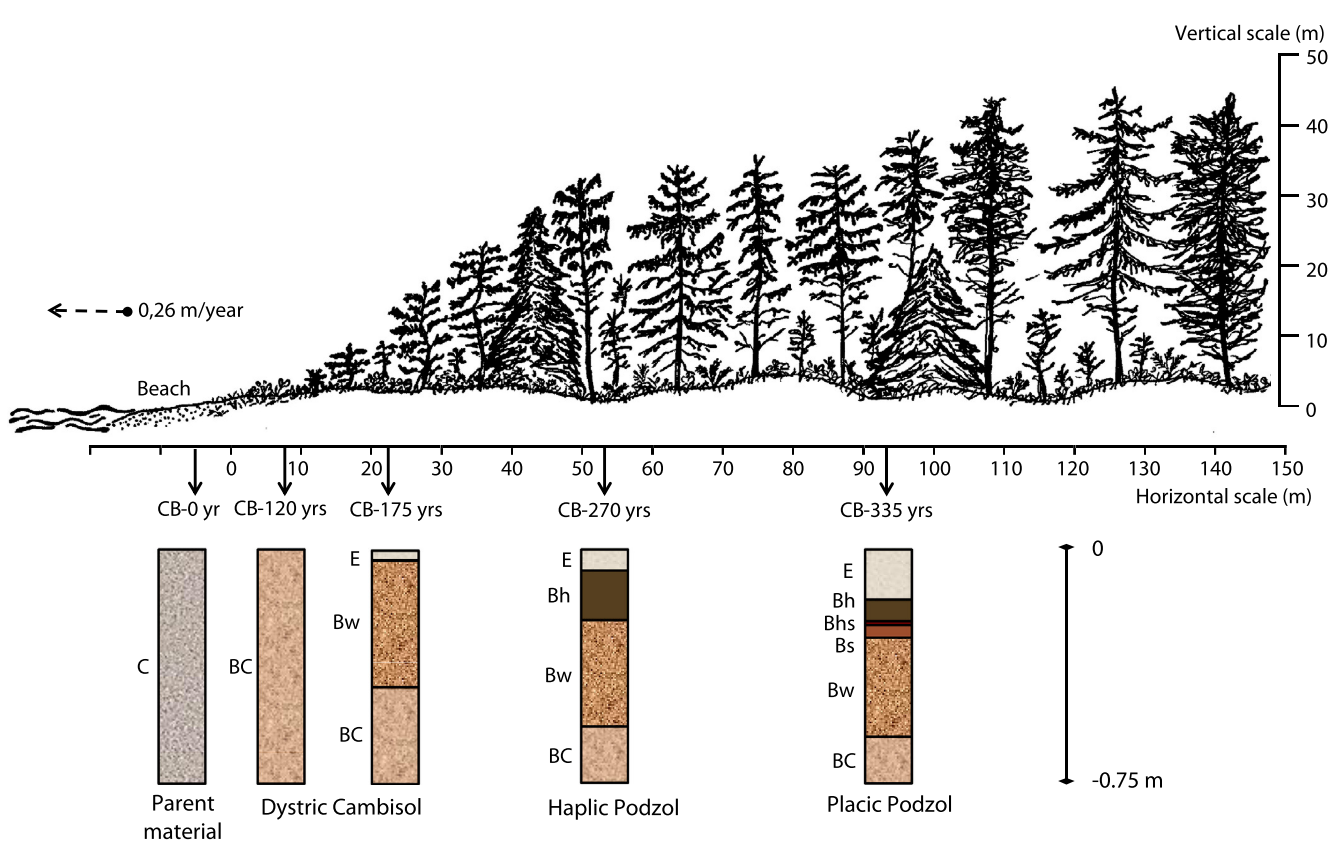

**BC**

**Ae**

**Bh**

**Supplementary material 3:** Details protocol of soil physico-chemical analysis

Briefly, soil pH was measured in 5g in 25 mL soil-solution suspension with CaCl_2_ and H_2_O. Cation exchange capacity and exchangeable cations were measured by ICP-OES following a 0.1N BaCl_2_ extraction. C% and N% were quantified using a FlashSmart Elemental Analyzer. Cold and hot C extraction were determined following the method described in (Ghani et al., 2003). Available ammonium (NH_4_) and nitrate (NO_3_) were assessed by 2N KCl extraction and auto-analyzer system. Organic P and inorganic P by ignition method followed by 0.5M H_2_SO_4_ (Saunders and William 1955, as modified by Walker and Adams 1958). Sodium pyrophosphate (0.1 M) and acidified ammonium oxalate (0.2 M) were used to extract Fe, Si, and Al from mineral samples, followed by Inductively Coupled Plasma Optical Emission Spectroscopy (ICP-OES) analysis (McKeague and Day, 1966; Courchesne and Turmel, 2008). Pyrophosphate-extractable Fe (Fe_pyr_) and Al (Al_pyr_) represent organically complexed Fe and Al, whereas oxalate extractable Fe (Fe_ox_) and Al (Al_ox_) represent amorphous and organically complexed Fe and Al. Sodium dithionite citrate extractable Al_dcb_, Fe_dcb_, Mn_dcb_ and Si_dcb_ comprised i) Al and Fe in organic complexes ii) Al and Fe in non-crystalline (hydr)oxides, and iii) Fe (and to a smaller extent the Al) in crystalline hydrous oxides. Mehlich III extraction followed by ICP-OES analysis was used to determine availability of Al, B, Ca, Cu, Fe, K, Mg, Mn, Na, P, S and Zn (Ziadi & Sen Tran, 2007). Total element concentration was measured by closed vessel microwave digestion with HNO3/HCl and analyzed by ICP-OES (U.S. Environmental Protection Agency, 1996).  Bulk soil mineral characterization by powder X-ray diffraction (XRD) was performed on a Phillips X’Pert Multipurpose X-ray diffractometer equipped with a fixed Cu anode operating at 45 kV and 40 mA. About 0.5 g of each soil was taken, homogenized by grinding in a pestle and mortar, and loaded into circular XRD sample holders. XRD patterns were collected in the 5–100^o^ 2*q* range with 0.1^o^ and 5s/step collection time. Analysis of the diffraction data for phase matching was carried out using JADE 9.5.1 (Materials Data, Inc.) and the PDF-4+ 2023 database from The International Centre for Diffraction Data (ICDD). Phase fractions for the minerals present were calculated from Rietveld refinement of the XRD data using the program TOPAS (Version 6, Bruker AXS, Germany) using full peak profiles.

**Supplementary material 4:** Soil physico-chemical properties. Data is presented as mean ± st-dev

|  | pH H_2_O | | | EC | | | C/N | | |
| --- | --- | --- | --- | --- | --- | --- | --- | --- | --- |
|  | (-) | | | (mS) | | | (-) | | |
| Ae | 4.46 | ± | 0.02 | 83.67 | ± | 0.72 | 9.5 | ± | 0.1 |
| Bh | 4.64 | ± | 0.02 | 32.63 | ± | 0.4 | 13 | ± | 0.1 |
| BC | 4.76 | ± | 0.01 | 91.73 | ± | 5.17 | 4.5 | ± | 0.4 |

| Cold C extraction | | | Hot C extraction | | | Available NH_4_ | | | | Available NO_3_ | | |
| --- | --- | --- | --- | --- | --- | --- | --- | --- | --- | --- | --- | --- |
| (mg/L) | | | (mg/L) | | | (mg/kg) | | | | (mg/kg) | | |
| 5.16 | ± | 0.76 | 28.25 | ± | 2.82 | | 5.7 | ± | 1.3 | 2.9 | ± | 0.2 |
| 4.84 | ± | 1.31 | 10.72 | ± | 1.55 | | 2 | ± | 0.3 | 0.2 | ± | 0 |
| 6.26 | ± | 1.34 | 18.66 | ± | 0.43 | | 1.4 | ± | 0.1 | 0.3 | ± | 0 |

|  | Exchangeable cations and CEC (cmol+/kg) | | | | | | | | | | | | | | | | | | | | | | | |
| --- | --- | --- | --- | --- | --- | --- | --- | --- | --- | --- | --- | --- | --- | --- | --- | --- | --- | --- | --- | --- | --- | --- | --- | --- |
|  | **Al** | | | **Ca** | | | **Fe** | | | **K** | | | **Mg** | | | **Mn** | | | **Na** | | | **Effective CEC** | | |
|  | (cmol+/kg) | | | (cmol+/kg) | | | (cmol+/kg) | | | (cmol+/kg) | | | (cmol+/kg) | | | (cmol+/kg) | | | (cmol+/kg) | | | (cmol+/kg) | | |
| Ae | 1.51 | ± | 0.09 | 0.41 | ± | 0.15 | 0.03 | ± | 0 | 0.06 | ± | 0.03 | 0.29 | ± | 0.15 | 0.01 | ± | 0 | 0.09 | ± | 0.02 | 3 | ± | 1 |
| Bh | 0.96 | ± | 0.48 | 0.27 | ± | 0.15 | 0.03 | ± | 0.01 | 0.04 | ± | 0.03 | 0.2 | ± | 0.16 | 0.01 | ± | 0 | 0.08 | ± | 0.05 | 1 | ± | 1 |
| BC | 0.49 | ± | 0.04 | 0.52 | ± | 0.04 | 0.04 | ± | 0.01 | 0.08 | ± | 0.01 | 0.45 | ± | 0.04 | 0.01 | ± | 0 | 0.16 | ± | 0.02 | 2 | ± | 0 |

| Texture, Hydrometer | | | |
| --- | --- | --- | --- |
|  | **Sand** | **Silt** | **Clay** |
|  | **%** | **%** | **%** |
| *Detection limit* | *2* | *2* | *2* |
| **Ae** | 82 | 14 | 4 |
| **Bh** | 91 | 6 | 3 |
| **BC** | 98 | <DL | <DL |

|  | Mehlich III extraction | | | | | | | | | | | | | | | | | |  |
| --- | --- | --- | --- | --- | --- | --- | --- | --- | --- | --- | --- | --- | --- | --- | --- | --- | --- | --- | --- |
|  | Al | | | Ca | | | Fe | | | K | | | Mg | | | Na | | | P |
|  | (mg/kg) | | | (mg/kg) | | | (mg/kg) | | | (mg/kg) | | | (mg/kg) | | | (mg/kg) | | | (mg/kg) |
| Ae | 782 | ± | 399 | 67 | ± | 21 | 27 | ± | 6 | 20 | ± | 10 | 37 | ± | 13 | 27 | ± | 6 | < DL |
| Bh | 1615 | ± | 1505 | 58 | ± | 59 | 30 | ± | 10 | 20 | ± | 10 | 29 | ± | 30 | 30 | ± | 10 | < DL |
| BC | 410 | ± | 86 | 112 | ± | 23 | 43 | ± | 6 | 40 | ± | 10 | 62 | ± | 3 | 43 | ± | 6 | 90 ± 10 |

**Supplementary material 5:** Relationship between total concentration of OA and root biomass, leaf biomass and area

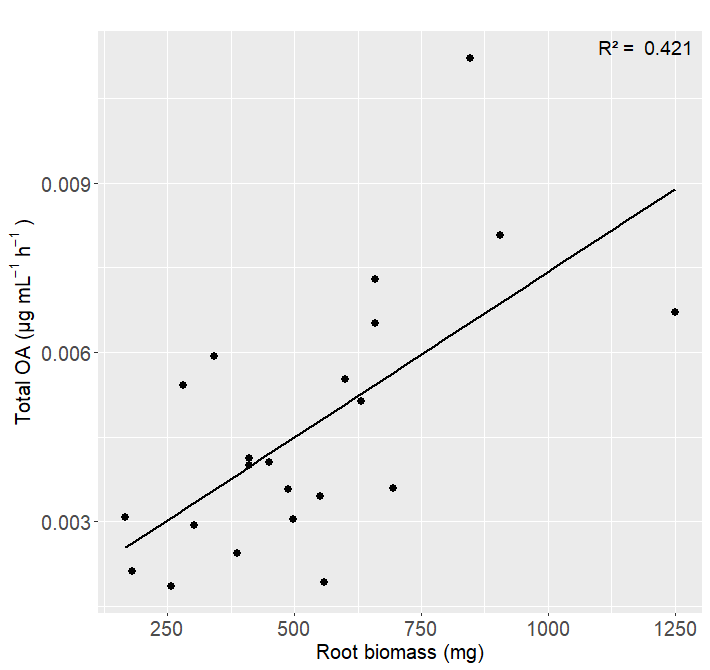

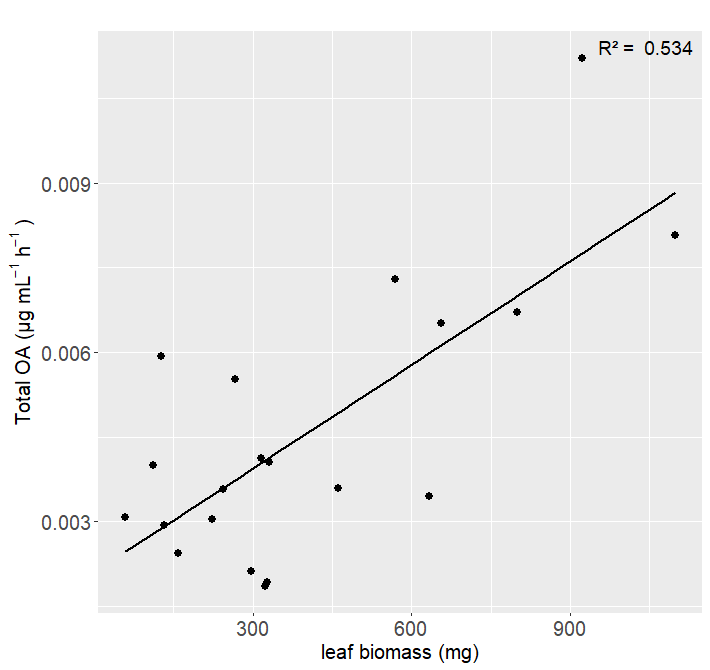

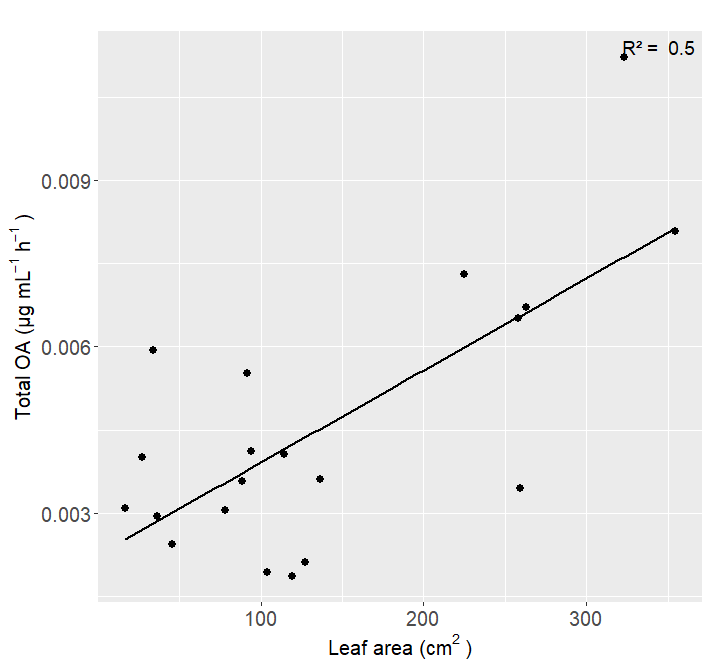

Supplementary material 6: Results from the 2 ways ANOVA assessing the effect of soil horizon, OA bin and their interactions on element dissolution. Different letters from the Tukey’s HSD post hoc test identify significant differences between group means.

| **Elements** | **Soil (pvalue)** | **OA_bin (pvalue)** | **Soil:OA_bin  (pvalue)** | **Post-hoc groups (OA_bin)** | **Post-hoc groups (Soil)** |
| --- | --- | --- | --- | --- | --- |
| **Mg** | 0.0114 * | 3.78e-10 *** | ns | High: a Medium: a Low: a Control: b | Ae: a BC: a Bf: b |
| **Al** | 2.23e-08 *** | 2.53e-05 *** | **0.00811 **** |  |  |
| **SI** | < 2e-16 *** | 2.44e-07 *** | ns | High: a Medium: ab  Low: b Control: c | Ae: b BC: b Bf: a |
| **K** | ns | 8.61e-08 *** | ns | High: a Medium: ab Low: b Control: c | **Ae:  BC:  Bf:** |
| **P** | 0.0279 * | 0.0102 * | ns | High: a Medium: ab Low: b Control:b | Ae: b BC: a Bf: c |
| **Ca** | ns | 3.84e-08 *** | ns | High: a Medium: a Low: a Control: b | **Ae:  BC:  Bf:** |
| **Fe** | 0.000555 *** | 2.11e-12 *** | **0.000110 ***** |  |  |

**Supplementary material 7:** Co-dissolution between 1) Al and Fe, 2) Si and Fe and, 3) Ca and Mg. Linear regression relationship between the elements across the three soil horizons. Tables summarize regression parameters including intercepts, slopes, standard-error, p-value, R2, adjusted R2 as well as the equations derived from the model by soil horizon.

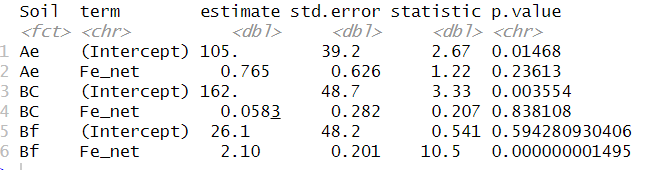

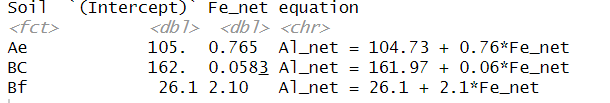

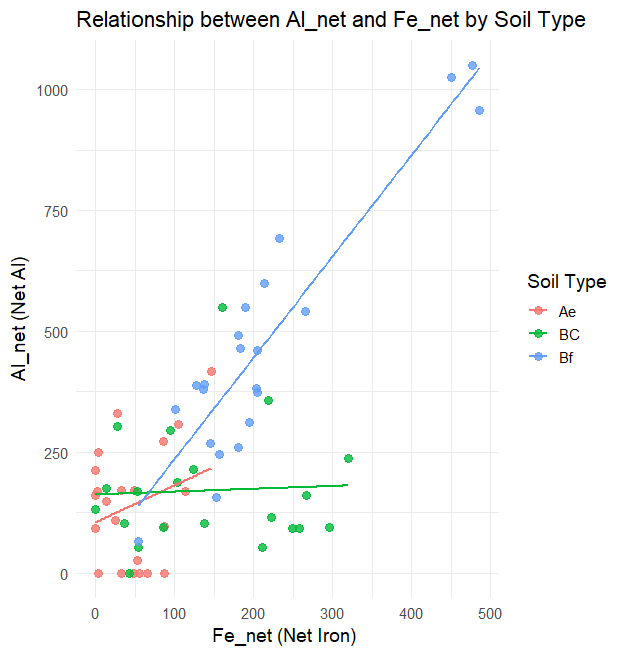

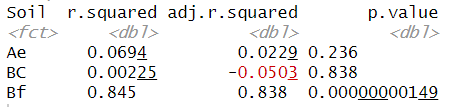

**1**

Bh

Bh

Bh

Bh

Bh

Bh

Bh

Bh

Bh

Bh

Bh

Bh

**2**

**3**

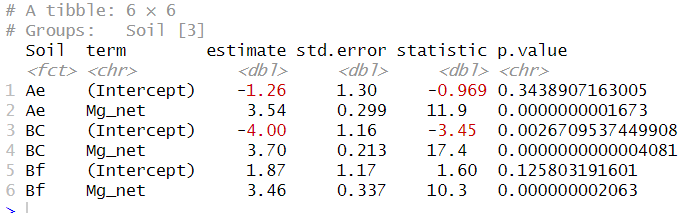

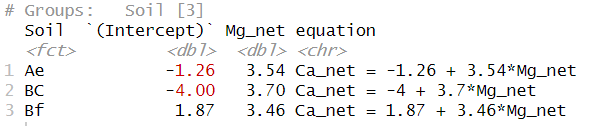

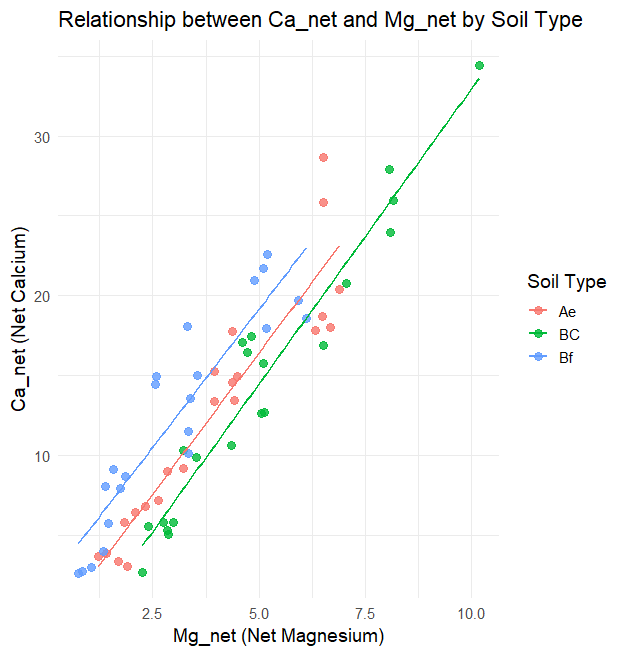

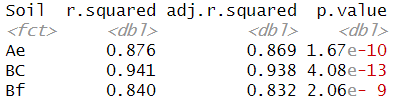

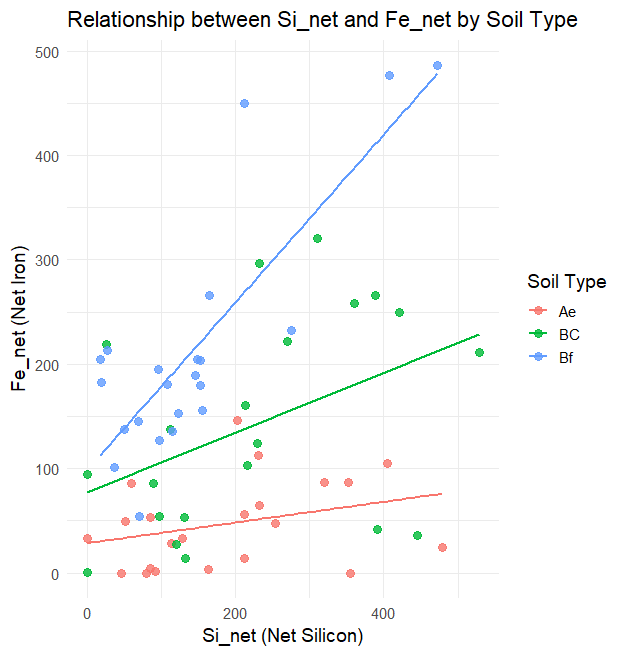

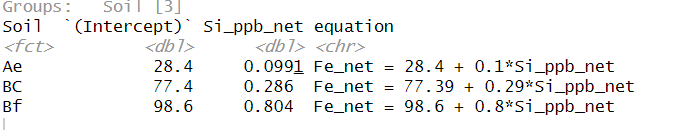

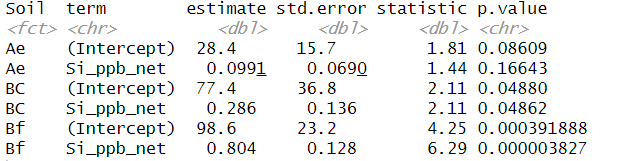

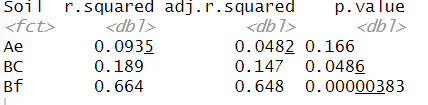
